# *rpfF* is not required for *X. translucens* pv. undulosa pathogenesis

**DOI:** 10.1101/2024.05.20.594922

**Authors:** Nathaniel Heiden, Jules Butchacas, Taylor Klass, Stephen P. Cohen, Veronica Roman-Reyna, Marcus V. Merfa, Guillermo E. Valero David, Christian Vargas-Garcia, Cristian Olmos, Yesenia Vélez-Negrón, Monica M. Lewandowski, Jonathan M. Jacobs

## Abstract

Bacterial cells self-coordinate via a mechanism called quorum sensing. In *Xanthomonas* species the gene *rpfF* encodes the quorum sensing autoinducer synthase. *Xanthomonas* species are divided into two main phylogenetic groups called Clade I and Clade II. The *rpf* quorum sensing system has been well studied in multiple Clade II *Xanthomonas* species and deletion of *rpfF* resulted in a major loss of virulence on susceptible hosts. However, the only Clade I *Xanthomonas* species in which the *rpfF* system was previously studied was in the sugarcane pathogen *X. albilineans*. In *X. albilineans* the *rpf* cluster plays a relatively small role in pathogenesis. *Xanthomonas translucens* pv. undulosa (Xtu) is a Clade I *Xanthomonas* species that causes bacterial leaf streak (BLS) and black chaff of wheat and barley and has increased as a concern in recent decades. Neither major resistance nor chemical treatments are available to prevent disease caused by Xtu. Interference with *rpf* bacterial quorum sensing systems has demonstrated some success in other systems. It was unknown whether BLS caused by Xtu could be prevented via quorum sensing interference. We found that Xtu encodes an *rpfF* homolog and we created an *rpfF* knockout mutant to study the role of the *rpf* system in Xtu. We found that the *rpfF* mutant was unaffected in its pathogenesis as it caused BLS symptoms and multiplied within wheat plants to the same levels as the wildtype strain. The Xtu *rpfF* mutant grew normally in lag and log phases *in vitro*, however it exhibited a shorter stationary phase and an early death phase in plant-derived media. The importance of RpfF in Xtu’s life cycle is unknown, though it appears to carry out a role in population stability. Our research determined that *rpfF* is not a major Xtu pathogenicity factor. Therefore, we do not recommend the targeting of the *rpf* quorum sensing system as a preventative treatment for BLS of wheat.

## Introduction

Parasitic bacteria frequently complete complex life cycles including drastic shifts in population behavior as a response to the environment and to the bacterial population size and status (Joshi et al. 2021). During these shifts, bacterial cells self-coordinate via a mechanism called quorum sensing to change gene expression and thus behavior in a cell density-dependent manner (González and Keshavan 2006; Tomasz 1965; Whiteley et al. 2017; Nealson et al. 1970). In quorum sensing, an autoinducer molecule unique to a set of bacteria is produced and perceived by the bacterial community. The amount of autoinducer perceived by a bacterium’s cognate receptor is a proxy for the quantity of like individuals. When a high level of autoinducer is present, a signaling cascade leads to changes in gene expression and a shift to high cell density behavior(s) (González and Keshavan 2006).

Phytopathogenic bacteria express diverse virulence factors during disease establishment and progression in their hosts (Benali et al. 2014; Pontes et al. 2020; Timilsina et al. 2020). Quorum sensing often regulates the expression of bacterial virulence factors, because many bacteria are required to express them simultaneously for efficacy (Joshi et al. 2021).

Xanthomonads, including phytopathogenic *Xanthomonas* species, use *cis*-2-unsaturated fatty acids called diffusible signal factors (DSFs) as a distinct family of autoinducers. Genes involved in *Xanthomonas* DSF production, sensing and response are located in the regulation of pathogenicity factors (*rpf*) cluster (Tang et al. 1991). DSFs are known to be produced via the fatty acid synthesis elongation cycle, with the thioesterase RpfF, encoded by *rpfF*, completing a crucial final step in the pathway (Zhou et al. 2015; Bi et al. 2014).

The loss of *rpfF* eliminates DSF production, preventing shifts in transcription related to DSF at high cell density (Figure 1). Therefore, *rpfF* deletion mutants have been used as a target to study the role of quorum sensing in *Xanthomonas* (Dow et al. 2003; Singh et al. 2022; Chatterjee and Sonti 2002; Rott et al. 2013; Huang et al. 2013). *Xanthomonas* species are divided into two main phylogenetic groups called Clade I and Clade II (Parkinson et al. 2007; Ferreira-Tonin et al. 2012; Jacques et al. 2016; Grau et al. 2016). The *rpf* quorum sensing system has been well studied in multiple Clade II *Xanthomonas* species. Deletion of *rpfF* resulted in a major loss of virulence on susceptible hosts for *X. campestris*, *X. oryzae*, *X. citri* and *X. axonopodis* (Dow et al. 2003; Chatterjee and Sonti 2002; Huang et al. 2013; Thowthampitak et al. 2008). DSF produced via RpfF was also found to have roles in motility and biofilm formation, however these varied depending on the species, and even strain, in question (Dow et al. 2003; Huang et al. 2013; Chatterjee and Sonti 2002). This fits with the broader Xanthomonadacae, as deletion of *rpfF* from *Xylella fastidiosa* resulted in hypervirulence *in planta* because *X. fastidiosa* has a drastically different lifestyle from characterized *Xanthomonas* species (Newman et al. 2004).

**Figure 1.**
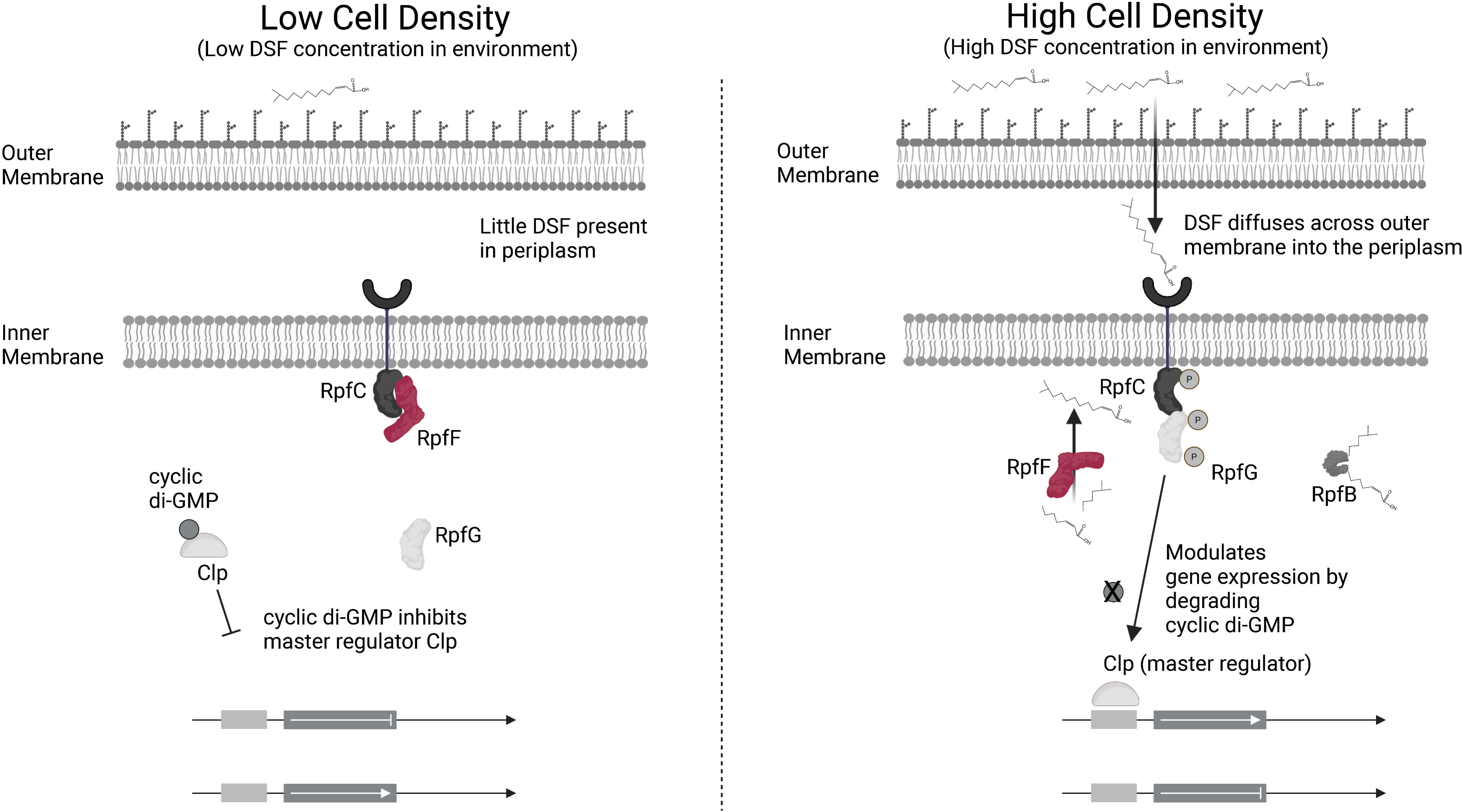
The regulation of pathogenicity factors (*rpf*) cluster in Xanthomonads. Proteins encoded by the *rpf* cluster mediate changes in gene expression in low cell density and high cell density. This figure was created with Biorender.com.

However, the only Clade I *Xanthomonas* species in which the *rpfF* system was previously studied was in the sugarcane pathogen *X. albilineans*. In *X. albilineans* the *rpf* cluster plays a relatively small role in pathogenesis (Rott et al. 2013). Virulence of an *rpfF* mutant was slightly, but significantly, decreased (Rott et al. 2013). Loss of *rpfF* also did not have an effect on *X. albilineans* motility (Rott et al. 2013). We do not know if the reduced importance of *rpfF* as the DSF synthase is unique to *X. albilineans* or if this is a characteristic of Clade I *Xanthomonas* species. *X. albilineans* also lacks the *hrp* regulon which is a link between the quorum sensing system and regulation of the Type III secretion system (T3SS) in *X. campestris* (Jiang et al. 2018; Pieretti et al. 2009). Other Clade I *Xanthomonas* species such as *X. translucens*, on the other hand, have an intact T3SS (Goettelmann et al. 2022).

*Xanthomonas translucens* pv. undulosa (Xtu) is a Clade I *Xanthomonas* species that causes bacterial leaf streak (BLS) and black chaff of wheat and barley (Sapkota et al. 2020). It can cause yield losses as great as 40% (Forster and Schaad 1988; Sapkota et al. 2020) and in recent decades has increased as a concern, especially in North America (Ledman et al. 2023; Tambong et al. 2021; Curland et al. 2018, 2020; Hangamaisho et al. 2024). Xtu is thought to spread by wind, rain and on seeds (Sapkota et al. 2020; Ledman et al. 2023). It enters leaves via natural openings such as stomata or damage points (Sapkota et al. 2020; Heiden et al. 2023; Adhikari et al. 2012). Once it gains entry to a wheat plant, Xtu extensively colonizes the apoplastic spaces of the mesophyll tissue (Heiden et al. 2023; Gluck-Thaler et al. 2020; Bragard et al. 1997; Sapkota et al. 2020; Jones et al. 1917). Once it reaches a high population, bacterial leaf streak manifests by macroscopic watersoaking symptoms followed by chlorotic and necrotic vein-delimited symptoms which give bacterial leaf streak its name (Ledman et al. 2023; Bamberg 1936; Jones et al. 1917; Sapkota et al. 2020). Once an Xtu population reaches a high level and encounters humid conditions, it oozes back out of stomata in exudates (Bamberg 1936). These exudates dry and are easily moved via wind or rain, enabling the spread to new leaves on the same or a different plant (Jones et al. 1917). In some cases, Xtu reaches the grain of a wheat plant and causes black chaff (Heiden et al. 2023; Bamberg 1936; Jones et al. 1917).

There is increasing interest in developing treatment options to combat BLS, including efforts to develop resistant germplasm in wheat (Sapkota et al. 2018), but currently no major resistance is available for growers. There are no available chemical treatments, and antibiotics can quickly drive the development of resistance when applied in the field (Sundin and Wang 2018). Plants broadly produce compounds which interfere with pathogen quorum sensing systems as part of their immune system (Joshi et al. 2021). Interference with bacterial quorum sensing systems has demonstrated some success in other systems. For example, plant expression of *rpfF* causes resistance to the Xanthomonad *Xylella fastidiosa* (Caserta et al. 2014, 2017; Lindow et al. 2014). However, it is currently unknown what role the *rpf* cluster plays in Xtu pathogenesis.

In this study, we investigated the role of the Xtu ortholog of *rpfF*, which encodes the critical DSF synthase in other *Xanthomonas* species. We tested the hypothesis that *rpfF* plays a similar major role in pathogenesis and plant colonization in Xtu. Specifically, we compare *X. translucens* to the well characterized Clade II *Xanthomonas* species: *X. campestris*. Our research focused on determining whether the *rpf* quorum sensing system is a main determinant of plant colonization in Xtu and therefore a desirable target for development of new strategies to prevent BLS caused by Xtu.

## Materials and Methods

### Bacterial strains and culture conditions

Bacterial strains were grown on nutrient agar (NA; beef extract 3g/L, peptone 5g/L, agar 15g/L), nutrient broth (NB; beef extract 3g/L, peptone 5g/L), Luria-Bertani (LB; 10g/L tryptone, 10g/L NaCl, 5g/L yeast extract) broth or Luria-Bertani agar (LBA; 10g/L tryptone, 10g/L NaCl, 5g/L yeast extract, 15g/L agar), or described media with supplemented 50µg/mL kanamycin, 40μg/mL spectinomycin, or 5μg/mL of gentamycin when applicable. Bacteria were grown for two or three days on NA plates at 28°C and then an overnight growth was resuspended to the desired optical density for the experiments in this study.

### Knockout mutant generation

To generate an in-frame deletion of *rpfF* in Xtu strain UPB513, genomic DNA was first extracted with the DNeasy Blood & Tissue Kit (QIAGEN, Cat. No. 69504) using TRIzol^TM^ (Invitrogen, catalog #15596026) following the manufacturer’s protocol with the following modification: 2mL of cell suspension with a concentration of 0.200 optical density at 600nm (O.D._600nm_) was centrifuged and resuspended with 0.75 mL TRIzol^TM^. The quality of genomic DNA was ensured via running in a 1.5% agarose gel at 110 V for 40 minutes. Upstream and downstream PCR fragments containing overhang regions with each other and the plasmid vector pk18*mobsacB* were amplified using the following two pairs of primers: rpfF-del-F1 and rpfF-del-R1 (Downstream fragment #1) and rpfF-del-F2 and rpfF-del-R2 (Upstream fragment #2) designed from the Xtu UPB513 genome (NCBI GenBank assembly GCA_023221635.1). The PCR products were cleaned using using the QIAquick® PCR Purification Kit (Qiagen, Cat. No. / ID: 28104) according to the manufacturer’s instructions. The vector pK18*mobsacB* (Schäfer et al. 1994) was linearized with HindIII and cleaned using the QIAquick® PCR Purification Kit. The two fragment products of *rpfF* and the linearized pK18*mobsacB* (Schäfer et al. 1994) were then isothermally assembled using the Gibson cloning kit (NEB; Gibson et al. 2009) following the manufacturer’s instructions in a 5:1 insert to backbone ratio. Primers rpfF-ko-conf-F and rpfF-ko-conf-R were used to confirm the construct.

*E. coli* RHO3 (López et al. 2009) cells were grown overnight in LB supplemented with N-succinyl-L-diaminopimelic acid desuccinylase and then subcultured for one hour until they reached an approximate 0.3 O.D._600nm_ in 50mL of LB broth. The suspension was centrifuged at 4000 x *g* for five minutes and then washed with 5mL of ice cold CaCl_2_ before being resuspended in ice cold 10% glycerol. 5µL of Gibson assembly product containing the assembled vector was added to 150uL of the competent *E. coli* and incubated on ice for 10 minutes and then placed into a 42°C water bath for 45 seconds. The tubes were returned to ice for 2 minutes and then 1mL of LB was added and the culture was placed in a shaking incubator for 1 hour. All of the transformation mixture was plated onto LB plates supplemented with kanamycin. Resistant colonies were selected and tested for the presence of the assembled fragments with primers rpfF-ko-conf-F and rpfF-ko-conf-R. The colonies were then tested with the primers rpfF-ko-conf-F and rpfF-ko-conf-R for the correct fragment insertion. A positive colony was then grown overnight, and the transformation plasmid was extracted with the QIAprep Spin Miniprep Kit (Cat. No. / ID: 27106X4).

Xtu UPB513 cells were grown for two days on NA plates and then transferred into nutrient broth (NB) for overnight growth. Competent cells were generated from an overnight culture that was subjected to 4 washes with 10% glycerol and centrifugation at 4000 x *g*. 400ng of the transformation plasmid was added to 100uL of competent UPB513 cells along with 0.75 µL of TypeOne™ Restriction Inhibitor (Epicenter) and incubated on ice for 2 minutes. Cells were electroporated at 2.5kV. The transformation mixture was incubated for 3 hours at 28°C in 1mL of NB medium with 250rpm shaking and then plated onto NA plates supplemented with kanamycin. Resistant colonies were picked and then plated onto both NA with kanamycin and NA with kanamycin and 10% sucrose. Colonies that were growth deficient in the presence of sucrose were pooled and grown overnight in NB before being plated on NA with 10% sucrose. Colonies that were no longer inhibited by sucrose were selected and plated on NA with kanamycin and 10% sucrose and NA with only 10% sucrose. Those colonies that were now sucrose insensitive and kanamycin sensitive were selected as potential mutants. Mutants were tested with PCR using the primers rpfF-del-F1 and rpdF-del-R2 and amplicons were sequenced with Sanger sequencing to confirm a clean deletion of 672 nucleotides in *rpfF* to create UPB513Δ*rpfF*.

### Knockout mutant complementation and introduction of gentamycin resistance

NEB Quickload polymerase and the primers rpfF-clone-F and rpfF-clone R were used to clone the entirety of the *rpfF* promoter, open reading frame and terminator region from UPB513 genomic DNA with regions overlapping into pUC18-miniTn*7*-T-Gm digested with HindIII. The cloned sequence was inserted into pUC18-miniTn*7*-T-Gm (Choi and Schweizer 2006) with Gibson assembly and then *E. coli* dH10b competent cells were transformed with the assembled plasmid using the previously described heat shock protocol. Cells were selected on NA with gentamycin and checked for the presence of *rpfF* with the primers rpfF-del-F1 and rpfF-del-R2. The plasmids were extracted with a Qiagen Miniprep kit and then Sanger sequenced using the primers rpfF-clone-F, rpfF-clone-R, walkprimer1, walkprimer2 and walkprimer3 to ensure that there were no mutations present in the inserted sequence. UPB513 and UPB513Δ*rpfF* were grown for two days on NA and then grown overnight in 10mL NB to an O.D._600nm_ of approximately 0.9. Competent cells were generated using the protocol from Choi et al. (2006) Cells were resuspended in 1mL 330*mM* sucrose at the final step. 100µL of the UPB513Δ*rpfF* competent cells were mixed with 250ng of the pUC18-miniTn*7*-T-Gm plasmid containing the cloned *rpfF* and 250ng of pTNS3. The mixtures were electroporated at 2.5kV and 1mL of NB was immediately added. After three hours of outgrowth at 28°C with 250rpm shaking the transformed cells were plated on NA supplemented with gentamycin. Colonies that had gained resistance to gentamycin were confirmed for the presence of both wildtype and mutant alleles of *rpfF* with the primers rpfF-del-F1 and rpfF-del-R2 and for genomic insertion at the *att*Tn*7* site (Choi et al. 2005) with primers insertion-conf-F and insertion-conf-R to validate the strain UPB513Δ*rpfF*::miniTn*7*T-Gm-*rpfF*. 100μL of the UPB513Δ*rpfF* and UPB513 competent cells were also each transformed with the same protocol using 250ng of pUC18-miniTn*7*-T-Gm and 250ng of pTNS3 to generate the strains UPB513Δ*rpfF*::miniTn*7*T-Gm and UPB513::miniTn*7*T-Gm.

### In vitro growth curve assay

Bacterial strains are grown from a starting concentration of 0.0001 O.D._600nm_. Each replicate was grown in 100uL in a 96 well plate with shaking at 567cpm for 48-96 hours at 28°C in a Biotek plate reader (Agilent, Santa Clara CA, USA). Measurements of O.D._600nm_ were taken every 30 minutes.

### Plant inoculations

Three cm sections of the 1^st^ leaf of 14-day-old wheat cv. Chinese Spring seedlings were infiltrated with a water mock or a 0.0001 O.D._600nm_ suspension of UPB513::miniTn*7*T-Gm or UPB513Δ*rpfF*::miniTn*7*T-Gm. Plants were grown in a growth chamber with a 12 hour photoperiod, 70% humidity and 25°C temperature. Pots were enclosed after inoculation for three days.

### In planta growth curves

Pictures were taken of four infiltrated leaf sections from the treatment groups and then a 1.5mm^2^ hole punch was taken from each infiltrated section. Hole punches were macerated and serially diluted. Serial dilutions were plated on NA with gentamycin and colony forming units were counted at 5 days. This experiment was repeated twice. Percentage of water soaking symptoms were determined for pictures taken at day six with ImageJ (Schneider et al. 2012).

### Mesophyll fluid extraction

The protocol from Roman-Reyna et al. (2024) for barley mesophyll apoplastic fluid (BMAF) extraction was adapted by removing the ribitol spiking step because metabolites were not quantified by normalization to ribitol.

## Results

### Xtu encodes a homolog of rpfF

The presence of *rpfF* is required for the pathogenesis of multiple *Xanthomonas* species. We therefore hypothesized that Xtu encodes an ortholog of *rpfF.* The amino acid sequence of *X. campestris* pv. *campestris* (NCBI accession 3M6M_A, Supplemental File 1) was used as a query with NCBI’s tblastn tool (Gerts et al. 2006) against the genome of Xtu UPB513 (NCBI GenBank assembly GCA_023221635.1) which returned a single significant alignment with 93% coverage and 53.5% identity (E-value = 2e^-96^, Max score = 308). This open reading frame was identified as the Xtu ortholog of *rpfF* (Supplemental File 2).

### rpfF is not required for Xtu growth and pathogenesis

Mutants in *rpfF* in other Xanthomonads have been critically impaired in their ability to colonize and cause disease in host plants. Therefore, we hypothesized that Xtu requires *rpfF* to coordinate behavior *in planta* and cause BLS symptoms. To challenge our hypothesis, we conduced multiple *in planta* experiments. The end stage of disease in which wheat leaf tissue is completely necrotic was achieved equally in plants infected with wildtype UPB513 and with UPB513Δ*rpfF* (Figure 2A). We observed exudates in both wildtype UPB513 and UPB513Δ*rpfF*. We hypothesized that although the end state of disease may be similar, it is possible that loss of *rpfF* impacts Xtu’s colonization of its wheat host in a minor way that our high-inoculum inoculation could not detect. To examine *in planta* growth, we tracked the population of both UPB513 and UPB513Δ*rpfF* over eight days. In two independent replicates we found that the populations of both UPB513 and UPB513Δ*rpfF* were equal every day (Figure 2B). We also measured the percentage of water-soaking in leaves at 6 days and did not find any differences between the mutant and wildtype infected leaves (Figure 2C). *Xanthomonas* motility is impacted in *rpfF* mutants in multiple Clade II *Xanthomonas* species (Chatterjee and Sonti 2002; Dow et al. 2003; Huang et al. 2013), but not in an *rpfF* mutant in the Clade I species *X. albilineans* (Rott et al. 2013). We hypothesized that *rpfF* is required for motility in Xtu. However, we found no evidence that loss of *rpfF* led to impaired motility (Figure 2D).

**Figure 2.**
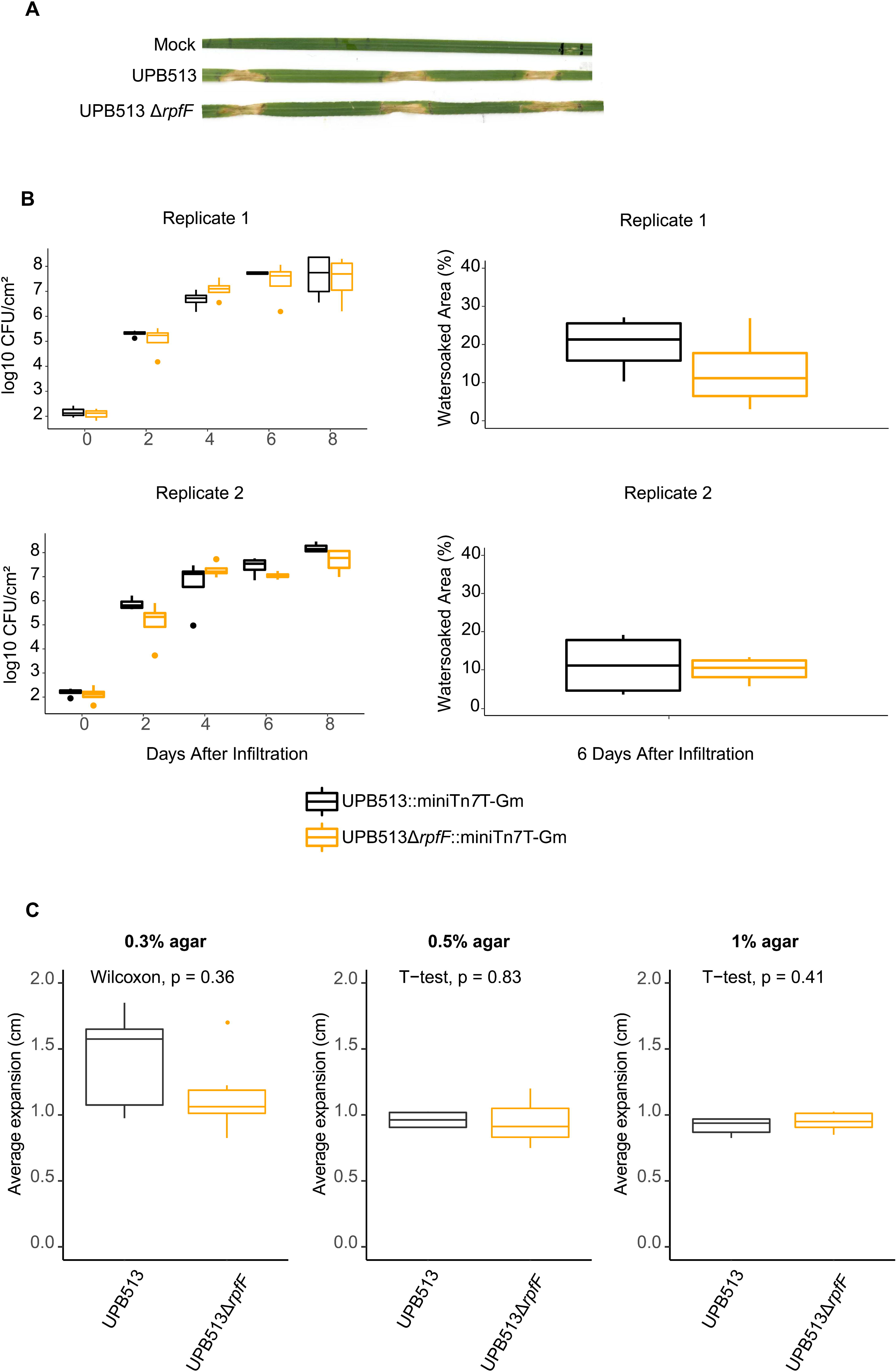
X. translucens does not require *rpfF* is not required for pathogenesis on wheat. (A) 3cm sections of the 1^st^ leaf of 14-day-old wheat cv. Chinese Spring seedlings were infiltrated with a water mock or a O.D._600nm_ 0.1 suspension of UPB513 and UPB513Δ*rpfF*. Plants were left covered for three days to create 100% relative humidity, and then were left uncovered for the subsequent 11 days. Exudates were observed in all treatments except for the mock at 3 days past inoculation (dpi). Pictures were taken at 14dpi. (B) *X. translucens* UPB513. 3cm sections of the 1^st^ leaf of 14-day-old wheat cv. Chinese Spring seedlings were infiltrated with a water mock or a O.D._600nm_ 0.0001 suspension of UPB513::miniTn*7*T-Gm, or UPB513Δ*rpfF*::miniTn*7*T-Gm. 1.5mm^2^ hole punches were taken from leaves of each treatment (n=4) at two, four, six and eight days after infiltration. Colony forming units were counted on selective media. At six days pictures were taken of samples leaves and percent water area was quantified with ImageJ. This experiment was repeated twice. (C) 5µL of UPB513 or UPB513Δ*rpfF* were spotted onto nutrient agar media plates with 0.3%, 0.5% or 1% agar. Colony expansion was measured after 21 days of growth at 28°C.

### rpfF is not required for normal growth in lag and log phases, but is required for maintenance of stationary phase in plant fluid

In *X. campestris*, *rpfF* plays important roles in life stage transition (Dow et al. 2003; Tang et al. 1991). We hypothesized that *rpfF* is required for normal growth and sustainment of Xtu in rich media. To test this, we compared the growth of UPB513 and UPB513Δ*rpfF in vitro* in both complex artificial media and fluid extracted from a plant environment. There were no differences in the lag, log and stationary phases between UPB513 and UPB513Δ*rpfF* in nutrient broth (Figure 3A). Growth rates were not significantly different between treatments. We found that UPB513Δ*rpfF* was also not deficient in lag and log growth phases in barley mesophyll apoplast fluid (BMAF) (Figure 3B). There was a clear difference in stationary phase in BMAF between UPB513 and UPB513Δ*rpfF*. UPB513Δ*rpfF* completed a normal shift to stationary phase, however a death phase began earlier in UPB513Δ*rpfF* than the wildtype UPB513 (Figure 3B). Wildtype stationary phase behavior was rescued by re-introduction via genomic insertion of the intact *rpfF* gene (Figure 3B). When run in the same *in vitro* experiments as previously described for Xtu, *X. campestris* Δ*rpfF* had irregular growth in lag and log phase in rich media (Figure 3C) but also exhibited a similar early death phase as demonstrated by *X. translucens* in BMAF (Figure 3D).

**Figure 3.**
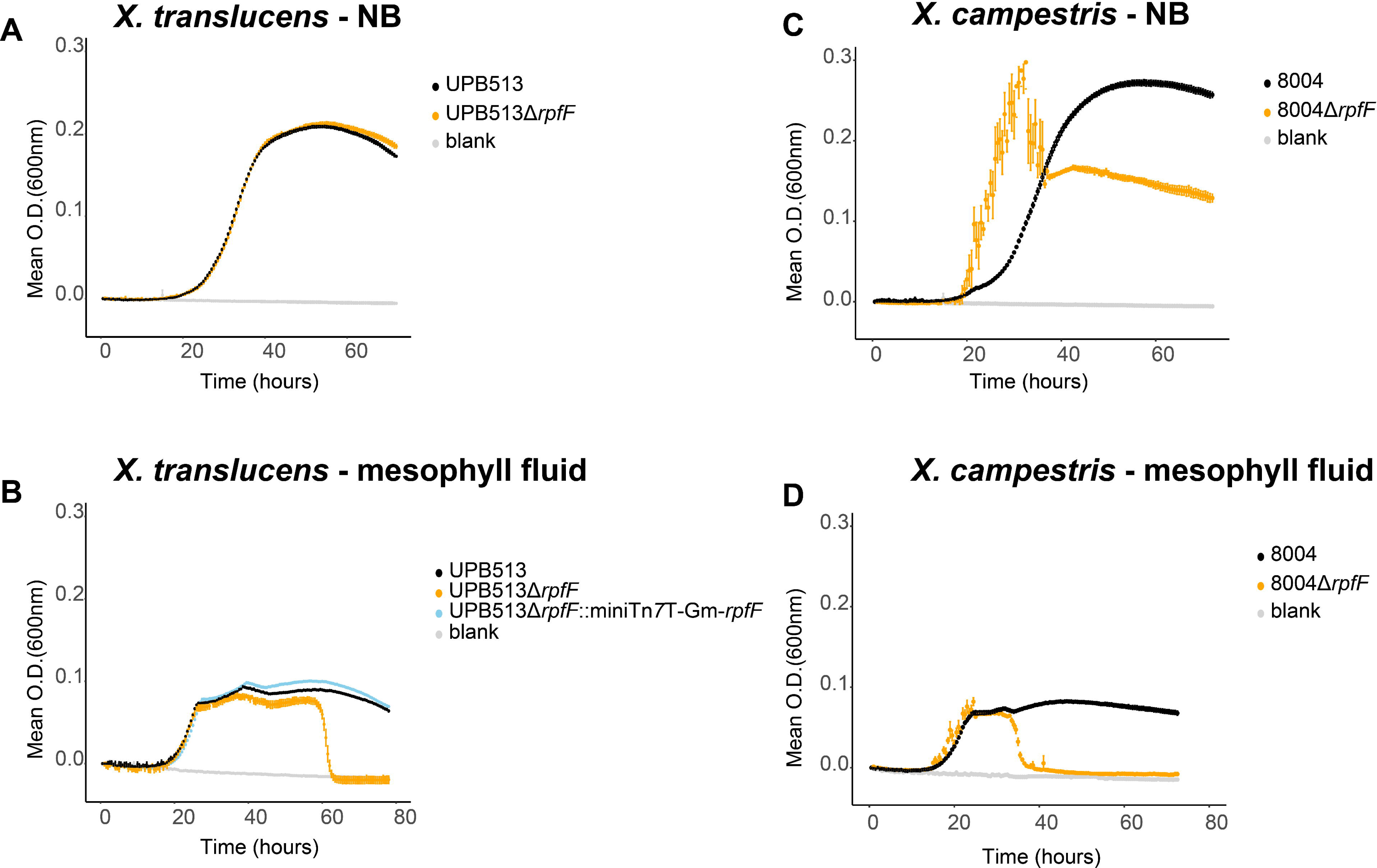
*rpfF* is required for normal *in vitro* growth of *X. translucens* and *X. campestris.* 4 replicates of each treatment were grown in 100uL in a 96 well plate with shaking at 567cpm for 72 hours at 28°C in a Biotek plate reader. A starting concentration of 0.0001 O.D._600nm_ of *X*. *translucens* UPB513 wildtype, UPB513Δ*rpfF*, UPB513Δ*rpfF*::mini-Tn7T-Gm-*rpfF* or a water control were inoculated into (A) nutrient broth or (B) barley mesophyll apoplast fluid (BMAF). A starting concentration of 0.0001 O.D._600nm_ of X. campestris 8004 wildtype, 8004Δ*rpfF*, or a water control were inoculated into (C) nutrient broth of (D) BMAF.

## Discussion

In this study we found that Xtu encodes a homolog of the DSF synthase gene *rpfF*. We discovered that Xtu was able to cause BLS symptoms and colonize wheat plants equally as well without *rpfF*. The *rpfF* mutant also retained wildtype motility. The Xtu *rpfF* mutant was unaffected in lag and log phases *in vitro* in both rich media and BMAF. The *X. campestris rpfF* mutant, in contrast, exhibited irregular growth in log phase in rich media. We found evidence that *rpfF* does play some role in stationary phase for both Xtu and *X. campestris* as population decrease occurred earlier for *rpfF* mutants.

The decreased importance of *rpfF* for Xtu may be a feature of Clade I *Xanthomonas* species. The gene *rpfF* has been well studied in multiple Clade II *Xanthomonas* species and found to play important roles in plant colonization and virulence in all cases. In stark contrast, Xtu colonized its primary niche, the mesophyll apoplastic space of wheat leaves, as normal despite deletion of *rpfF*. We speculate that some evolutionary pressures, such as plant recognition of DSF, may have led to changes in its function in Xtu. A *rpfF* mutant in *X. albilineans*, another Clade I *Xanthomonas* species, did have a decreased ability to colonize sugarcane stalks but this phenotype was much less extreme than what has been described in Clade II *Xanthomonas* species. Examination of the importance of this cluster in other Clade I species will be required to determine whether this pattern holds true.

*X. albilineans* and Xtu have substantially different lifestyles. Xtu is a nonvascular pathogen of cereal plants (Gluck-Thaler et al. 2020), while *X. albilineans* colonizes the vascular xylem of sugarcane plants (Rott et al. 2013). It is possible that Clade I *rpfF* may be relevant to vascular colonization, but not for nonvascular colonization such as that carried out by Xtu. Future work examining the role of *rpfF* of vascular *X. translucens* strains such as those in the pathovars translucens and graminis would test this hypothesis. *X. albilineans* also lacks the *hrp* regulon which is a link between the quorum sensing system and regulation of the T3SS in *X. campestris* (Jiang et al. 2018; Pieretti et al. 2009). Xtu on the other hand, has an intact T3SS (Goettelmann et al. 2022). Therefore, it’s probable that *X. albilineans* and Xtu have evolved distinct strategies to utilize and regulate their T3SS during plant colonization suggesting that the function of rpfF may not be analogous in both cases.

We described a role for *rpfF* in stationary phase in both Xtu and *X. campestris*. In summary, Xtu was able to grow normally without *rpfF*, however in late stationary phase the optical density, which is a proxy for population, decreased. There was a similar population decrease for both Xtu and *X. campestris*, however this effect was more pronounced and occurred earlier for *X. campestris* which correlates with the more important role that *rpfF* plays in *X. campestris.* Interestingly, this only occurred in plant-extracted mesophyll fluid, and not in artificial rich media.

In *X. oryzae*, Singh et al. (2022) determined that *rpfF* plays a role in maintaining membrane stability. The stress experienced by a cell increases in stationary phase relative to log phase (Singh et al. 2022). It is unknown whether there is a link between membrane stability and the phenotype we saw in the stationary phase of UPB513Δ*rpfF*. The existence of this death phase phenotype in an Xtu *rpfF* mutant suggests that *rpfF* does indeed play some role in coordinating cell-to-cell communication in Xtu populations.

The high cell density switch to stationary phase *in vitro* and to virulence-related behaviors *in planta* is not fully explained by the *rpf* system in Xtu. This strongly suggests that additional signaling systems are involved in perception of and response to high cell density. We cannot rule out the additional possibility that this system is still responsible for perception of an autoinducer signal via RpfC, and that *rpfF* is not required for autoinducer synthesis. The in-vitro loss of planktonic growth phenotype we discovered could be used in a high-throughput screening of mutants to find additional genes important for quorum sensing.

A main question that remains as a result of our research is what role does *rpfF* play in Xtu’s life cycle. Xtu encodes *rpfF* which implies a selective pressure to retain this gene and loss of *rpfF* results in changes in stationary phase behavior *in vitro*. However, there was no discernible importance of *rpfF* in a susceptible wheat plant. Xtu has a wide host range (Heiden et al. 2023) and *rpfF* may be required for colonization of other hosts or more resistant wheat varieties which present more of a challenge for Xtu. It is also possible that this gene is important for environmental survival in suboptimum nutrient environments.

Our main objective was to determine whether *rpfF* was a critical pathogenicity factor that could be interfered with to combat BLS of wheat. Our research determined that *rpfF* is not a major Xtu pathogenicity factor. Therefore, we do not recommend the targeting of the *rpf* quorum sensing system as a preventative treatment for BLS of wheat.

**Table 1.**
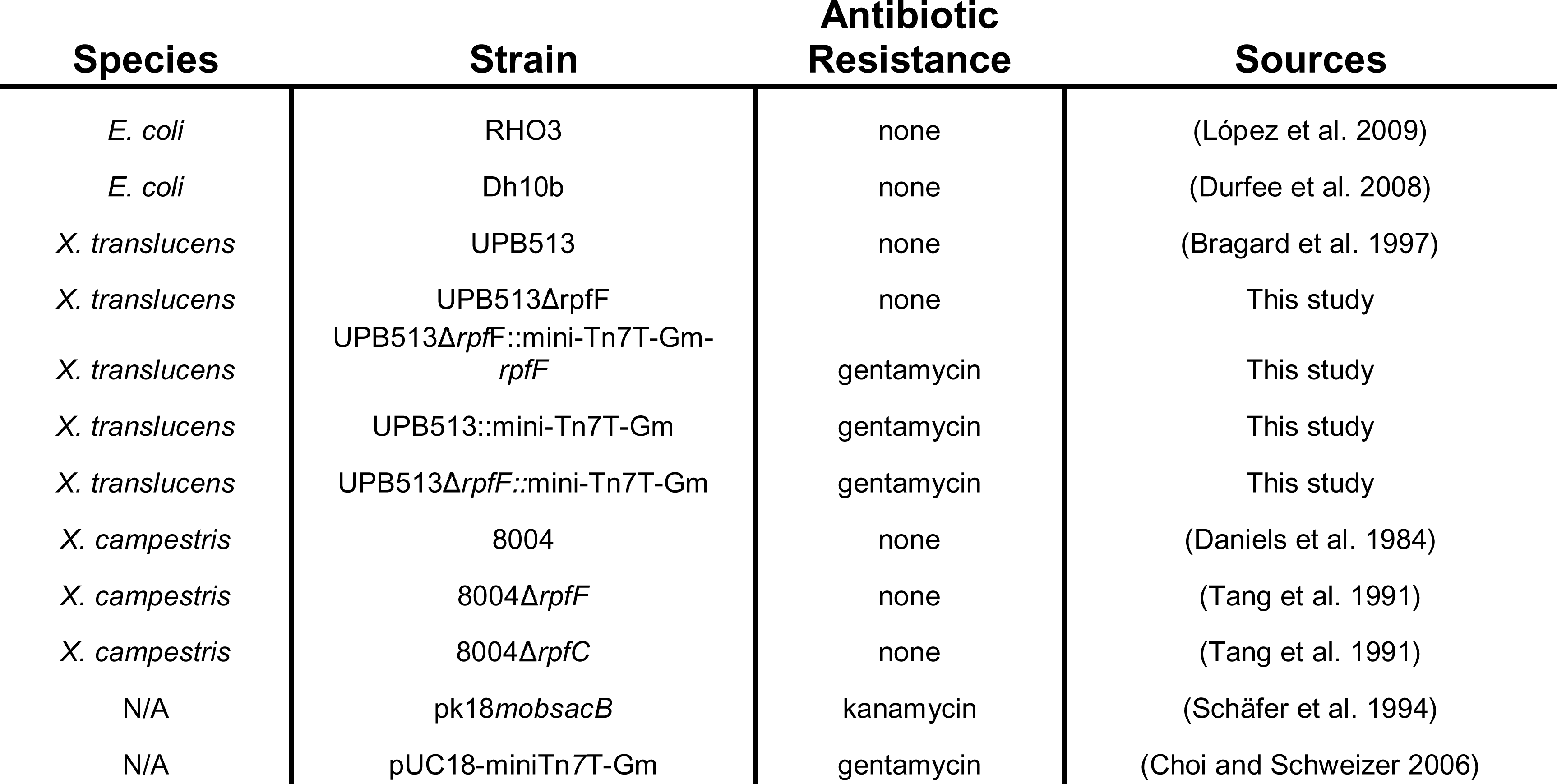
Bacterial strains and plasmids used in this study.

| Species | Strain | Antibiotic Resistance | Sources |
| --- | --- | --- | --- |
| *E. coli* | RHO3 | none | (López et al. 2009) |
| *E. coli* | Dh10b | none | (Durfee et al. 2008) |
| *X. translucens* | UPB513 | none | (Bragard et al. 1997) |
| *X. translucens* | UPB513Δ*rpfF* | none | This study |
| *X. translucens* | UPB513Δ*rpfF*::mini-Tn7T-Gm-*rpfF* | gentamycin | This study |
| *X. translucens* | UPB513::mini-Tn7T-Gm | gentamycin | This study |
| *X. translucens* | UPB513Δ*rpfF*::mini-Tn7T-Gm | gentamycin | This study |
| *X. campestris* | 8004 | none | (Daniels et al. 1984) |
| *X. campestris* | 8004Δ*rpfF* | none | (Tang et al. 1991) |
| *X. campestris* | 8004Δ*rpfC* | none | (Tang et al. 1991) |
| N/A | pk18*mobsacB* | kanamycin | (Schäfer et al. 1994) |
| N/A | pUC18-miniTn7T-Gm | gentamycin | (Choi and Schweizer 2006) |

**Table 2.**
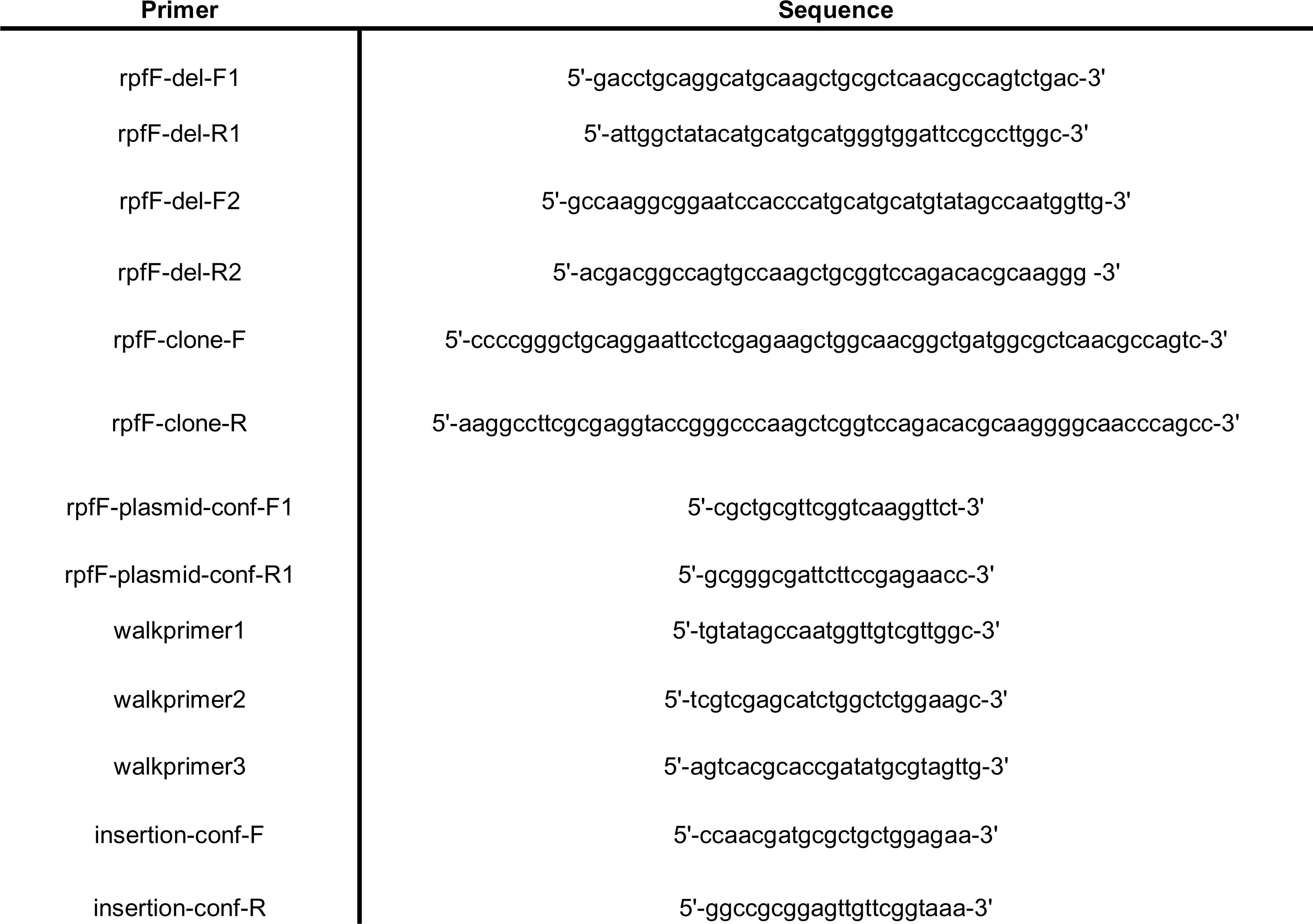
Primers used in this study.

## Acknowledgements

We thank Dr. Tuan Minh Tran for sending us the *X. campestris* strains used in the experiment and for helpful input during the preparation of this manuscript. We thank Zihang Gao for help running PCR assays.

## Authors disclaimer

Mention of trade names or commercial products in this report is solely for the purpose of providing specific information and does not imply recommendation or endorsement by the U.S. Department of Agriculture (USDA). USDA is an equal opportunity lender, provider, and employer.

---

Supplemental Figure 1. Knockout of *rpfF* gene in *X. translucens* UPB513. (A) A construct was designed to knockout out 224 amino acids of *rpfF*. (B) The deletion of 672 nucleotides in the *rpfF* gene, corresponding to 224 amino acids was confirmed by PCR amplification of the sequence of interest in the wild type (expected length 1223 base pairs) and mutant (expected length 544bp) strains and gel electrophoresis.

