## Supplementary figures and images for "*rpfF* is not required for *X. translucens* pv. undulosa pathogenesis"

### Supplemental Figure 1

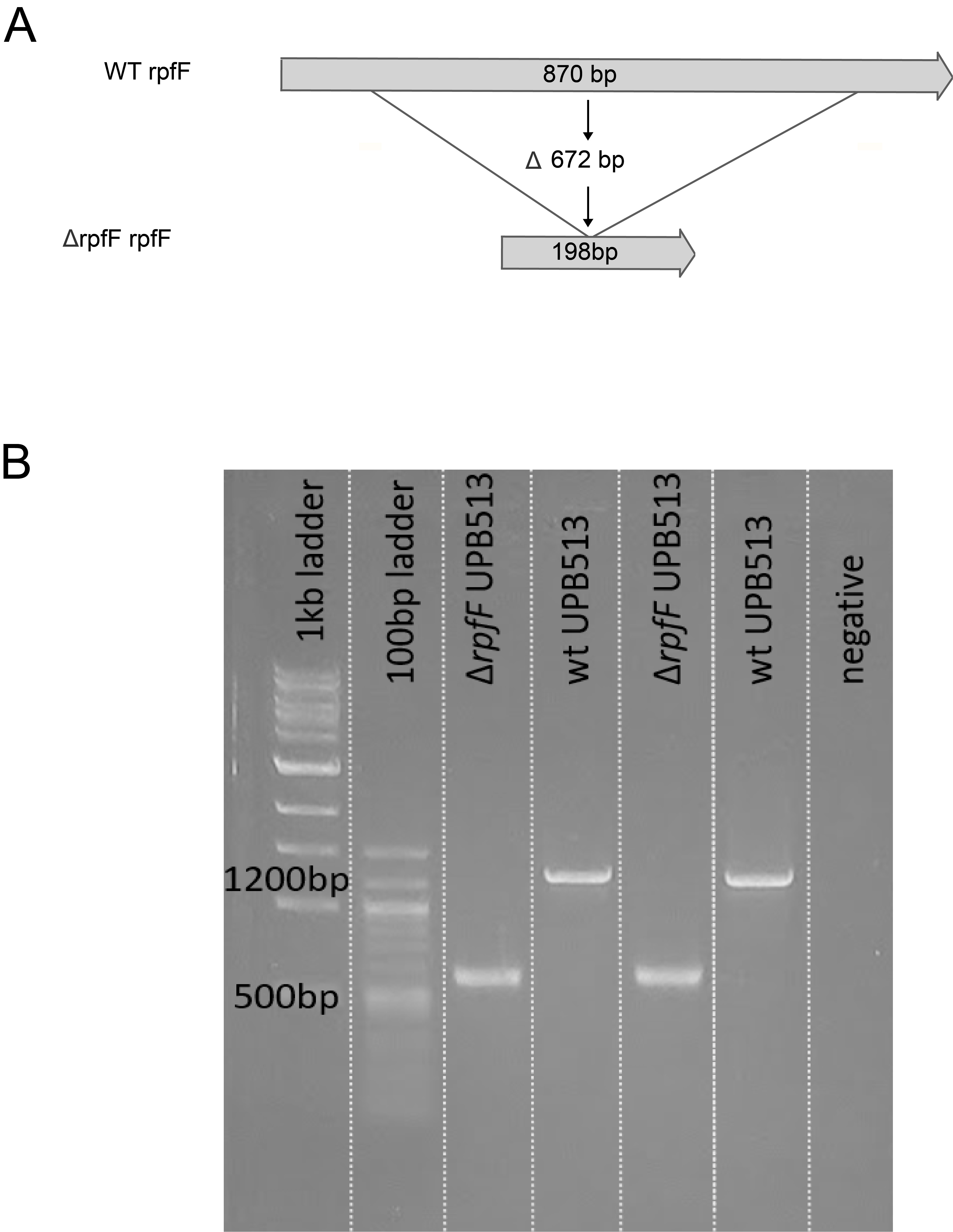
